# Interaction of Environment and Vineyard Management Effects Shape Microbial Functionality, *Terroir*, and Grape Qualities

**DOI:** 10.1101/2025.08.18.670824

**Authors:** Lena Flörl, Maria Paula Rueda-Mejia, Jean-Sébastien Reynard, Vivian Zufferey, Kathleen Mackie-Haas, Emmanuel Frossard, Nicholas A. Bokulich

**Author notes:** Corresponding author: Nicholas A. Bokulich. Address: ETH Zürich, LFW E 58.1, Universitätstrasse 2, 8092 Zürich,Switzerland.

## Abstract

Vineyards are complex agroecological systems, with geographically distinct microbial communities playing a central role in mediating vine health, grape quality, and ultimately local wine characteristics (*terroir)*. However, the extent to which environmental factors and vineyard management practices interact to shape these microbial communities and their functions remains poorly understood. We implemented a multidisciplinary citizen science collaboration together with winegrowers to sample 19 Pinot Noir vineyards across two distinct Swiss regions (N=38 samples). This allowed us to evaluate how a combination of environmental conditions and management practices (such as cover cropping and fungicide use) collectively impact soil and grape microbiomes, soil enzymatic activities, nutrient concentration in leaves, and grape chemistry. Regional differences were the primary drivers of variation, but multivariate analysis revealed a distinct, secondary influence from management practices on soil and berries. Berry quality was linked to functional soil properties, including microbial activity, which were further associated with key fermentative yeasts likely involved in wine fermentation. These findings highlight the interconnected nature of environmental, microbial, and agronomic factors in grapevine systems and offer novel insights into functional aspects of *microbial terroir*.

## Introduction

Understanding the factors that shape microbial communities and their function in agricultural ecosystems is relevant for answering fundamental ecological questions as well as in developing sustainable farming practices. Viticulture is one of the most agriculturally intensive systems with high inputs of pesticides and frequent reliance on irrigation. For example, in Europe, in 2007 viticulture occupied only 3.5% of the agricultural land but accounted for approximately two-thirds of all fungicide use (*The Use of Plant Protection Products in the European Union* 2007). This highlights the need for more sustainable approaches, in which microorganisms and their management could be a central factor (Trivedi *et al*. 2021).

Grapevine-associated microbial communities play critical roles in vine health, fruit development, and ultimately, wine quality (Zarraonaindia *et al*. 2015; Griggs *et al*. 2021). These communities are influenced by a complex interplay of biotic and abiotic factors, which include spatial, temporal and management-related variables (Griggs *et al*. 2021, 2025; Bettenfeld *et al*. 2022). Regional variation in these factors shapes microbial biogeography across space and time and lead to predictive associations between grapevine microbiota and wine qualities, a concept termed *microbial terroir* (Bokulich *et al*. 2014, 2016; Flörl *et al*. 2025c). However, the complexity of abiotic and biotic factors in commercial vineyard settings makes it challenging to unravel relative contributions of individual drivers.

As in every agricultural setting, soil is a central component of the vineyard ecosystem as it provides the vine with nutrients and water, and is a hotbed of microbial activities that influence nutrient uptake by the plant and growth characteristics. Nitrogen (nitrate and ammonium), sulfate and to a lesser extent phosphate in our soils are made available to the plant through microbial organic matter decomposition (Burns *et al*. 2013). Soil microbial communities, thereby, have a reciprocal relationship with soil properties, as their composition is influenced by their respective environmental conditions, while their function in turn actively modulates these (Philippot *et al*. 2024). In particular, extracellular soil enzymes – which are key to carbon, nitrogen, and phosphorus cycling – are highly sensitive to environmental conditions and vineyard practices (Giffard *et al*. 2022). Although their potential to indicate soil function and influence grape quality is increasingly recognized (Griesser *et al*. 2022), their specific role and interconnectedness with other factors in viticultural systems has not been widely explored. The complexity of such interactions calls for interdisciplinary and integrative studies (Giffard *et al*. 2022; Philippot *et al*. 2024). Yet, to control for some of these variables requires extensive and targeted recruiting, which is a common bottleneck when studying commercial viticulture.

To address this issue and to bridge the gap between fundamental research and practical needs in viticulture, we launched an interdisciplinary citizen science initiative in collaboration with commercial winegrowers and wine industry stakeholders. In workshops with various stakeholders, we co-developed relevant research questions and a targeted questionnaire to better understand growers’ perspectives, priorities, and questions. Based on this consensus, we investigated how microbial communities in soil and berries are interconnected with soil enzymatic activities and physicochemical properties, under varying environmental conditions and management practices, while accounting for grapevine cultivar. Based on the questionnaire responses, we selected representative vineyard sites from two climatically and pedologically distinct regions, all planted with Pinot Noir to control for cultivar/genotype-related effects (Flörl *et al*. 2025a), but with different cover crops usage, irrigation and cropping systems (see Figure 1). Alongside microbiome analyses, we collected comprehensive data on soil properties, nutrient concentration in leaves, and vineyard management practices. This integrated dataset enables us to identify key environmental and management factors shaping microbial community structure and function.

**Figure 1.**
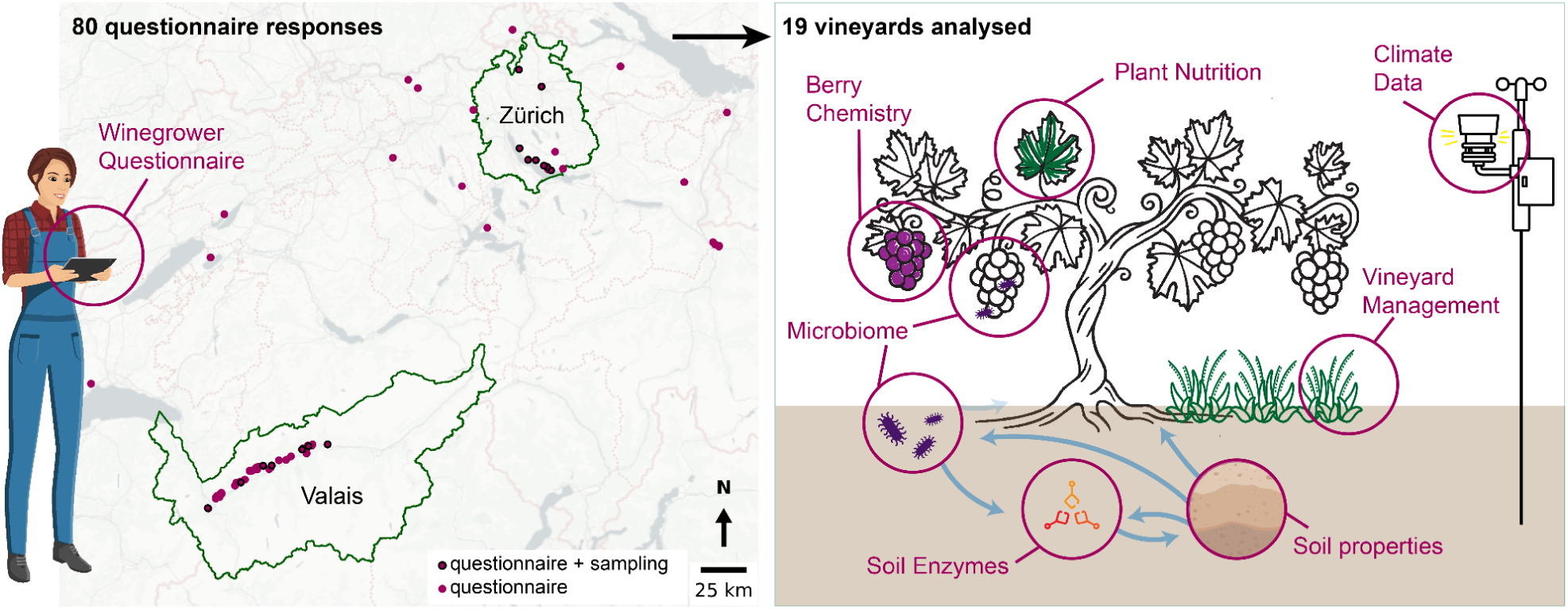
**Graphical Abstract**: We surveyed 80 Swiss winegrowers (purple circles) and subsequently analysed 19 vineyards (circles with black edge) from two climatically distinct regions (Valais and Zürich). At each site we collected berry and soil samples for microbiome profiling, as well as analysing soil properties, berry chemistry, and leaf nutrient status. Additionally, we collected local climate data and vineyard management practices. Our goal was to study the influencing factors shaping microbial communities and their function, specifically regarding soil enzymatic activities.

## Results

### Integrating Stakeholders throughout the Research Process to Understand Management Choices and their Effects

The collection of samples from vineyards is a bottleneck in many research endeavors and a key reason why close collaboration with winegrowers is central to evaluate *microbial terroir* effects on vineyard settings. To address this challenge and to involve winegrowers directly, we launched a citizen science project to identify mutual aims for examining *microbial terroir* in the context of different viticultural management practices (see Supplementary Methods). Through co-design workshops with stakeholders, we identified key research priorities, like the role of microbial communities and sustainable management practices. Based on this input, we developed a questionnaire to understand winegrowers’ current practices and needs, gathering 80 responses from across Switzerland that offered several valuable insights into their motivations and perspectives (Figure 2 and Supplementary Figure 1). Consumer appreciation, suitability of the location, and adaptability to climate change are reported as primary motivations for selecting new grape varieties, but not financial incentives (Figure 2A). While soil type was widely regarded as having a strong influence on wine quality, the effect of location on disease occurrence was perceived as more moderate (Figure 2C)

**Figure 2.**
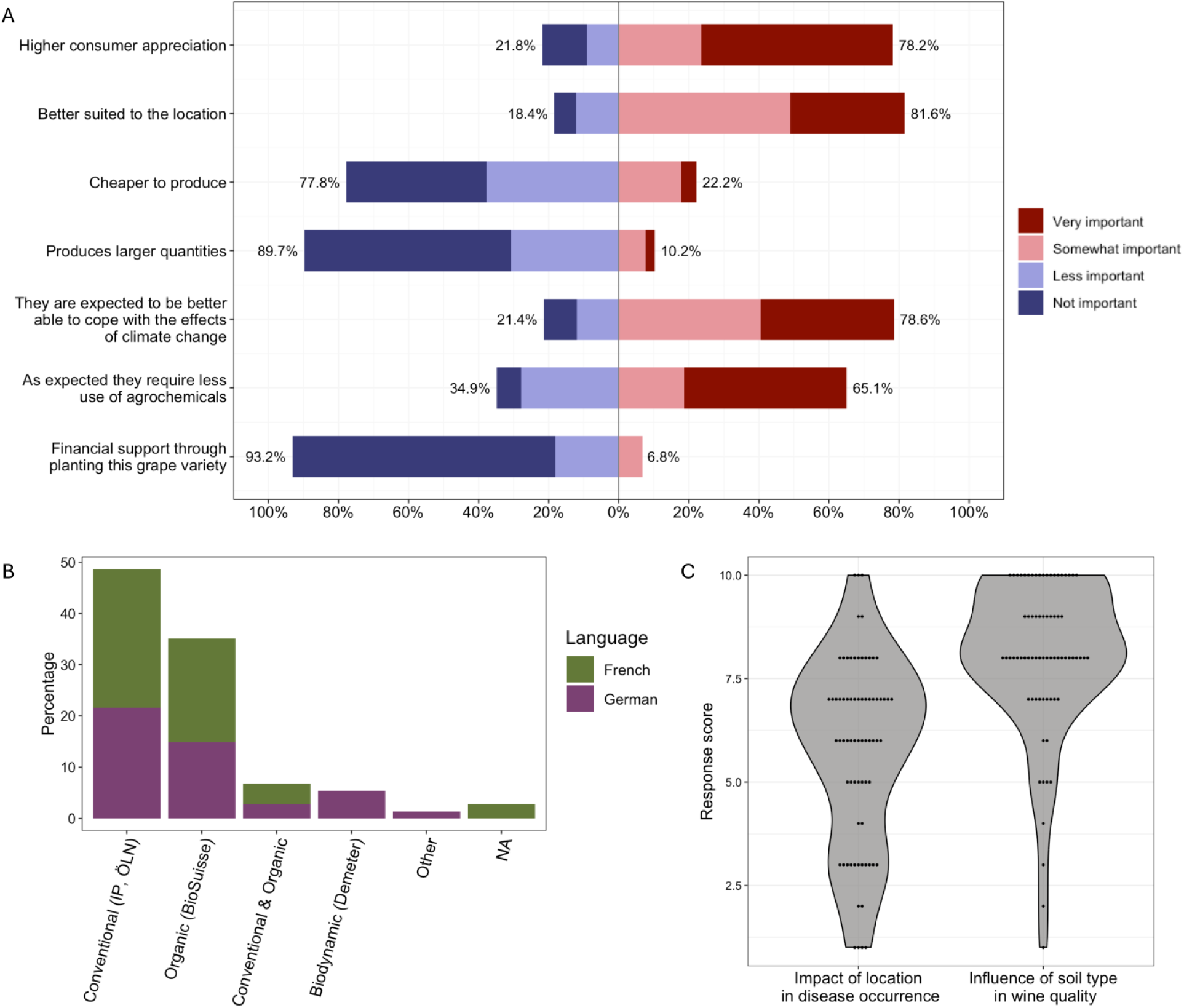
Insights on viticultural practices and perceptions among winegrowers. (A) The reasons for planting new varieties are mostly driven by anticipated consumer appreciation and local suitability, and to a lesser degree by direct financial incentives or yield. (B) The distribution of cropping systems are similar between regions. (C) Surveyed winegrowers perceive soil type to be important for wine quality (Response score: 10 being very important, and 1 being not important at all), while perceptions of whether disease occurrence varies by location are more diverse.

Most respondents practice conventional viticulture (49%), followed by organic production (35%), while 6.7% use either approach for different parcels of their production. Notably, biodynamic agriculture, which employs the principles of organic agriculture along with unique supplementary practices, as certified with the Demeter label (Castellini and and Troiano 2017), is practiced by 11.7% of the respondents from German-speaking regions (5.4% of the total). However, it is not used by any of the producers surveyed in francophone Switzerland (Figure 2B). Finally, most producers (69%) use their harvest to make wine themselves, while 20% sell their grapes (all in French-speaking regions), and a few (7%) outsource wine production (Supplementary Figure 1A).

### Microbial Communities of Vineyard Soils and Berries vary between Regions

Based on the questionnaire responses from winegrowers across Switzerland, we selected 19 vineyards located in the climatically and topographically distinct regions of canton Valais and canton Zürich. To parse biotic effects we exclusively sampled Pinot Noir vineyards to reduce variability associated with grapevine variety (Flörl *et al*. 2025b). Within each region, we identified representative plots reflecting a range of management practices, particularly under-vine floor management (cover crop usage), irrigation regimes, and cropping systems (see Supplementary Table 1). At these sites, we conducted comprehensive sampling to assess the soil and berry microbiomes (N = 38), alongside measurements of soil properties, berry chemistry, leaf nutrient content, and local climate conditions. While we controlled for cultivar and considered the context of various management practices, vineyard sites differed considerably in soil characteristics and climatic conditions between regions. Vineyard elevations ranged from 510 to 844 m in Valais and from 422 to 534 m in Zürich. The two regions also exhibit pronounced climatic differences: Zürich, located on the Swiss Plateau, experiences a more humid climate, whereas Valais, situated in the Swiss Western Alps, is among the driest regions in Switzerland. Sampled vineyards, therefore, vary significantly in annual rainfall (Valais: mean = 713 mm; Zürich: mean = 1171 mm; ANOVA: p=7.642e-18, R^2^=0.875) between the regions, as well as in average measured soil temperatures (Valais: mean = 13.80°C; Zürich: mean = 13.02°C; ANOVA: p=0.036, R^2^=0.116), but not in average air temperature (Valais: mean = 12.02°C; Zürich: mean = 11.90°C; ANOVA: p=0.224, R^2^=0.041). Additionally, soil types varied between regions: soils from the canton of Zurich were predominantly Cambisols (IUSS Working Group 2014), whereas those from canton Valais were crystalline, non-calcareous, sometimes slightly acidic scree (Canton du Valais 2023).

To study microbial communities in berries and soil in these different contexts, we performed amplicon marker gene sequencing (bacterial 16S rRNA gene and fungal ITS domain sequencing), yielding 537’405 and 823’927 reads for bacteria and fungi, respectively. Notably, due to insufficient sequencing depth for bacterial communities in berry samples, only fungal communities of berries were analyzed. While alpha diversities did not significantly vary between regions or management regimes in soil (Supplementary Table 2) and berries (Supplementary Table 3), community structures were markedly different between vineyards from different cantons (Figure 3). Notably, taking the abundance of taxa into account (Bray Curtis metrics) increases the variance explained by region, as does considering the phylogenetic relatedness of features (k-mer based metrics). In soil communities, the different regions thereby explained up to 36.8% of variance in bacteria and 21.8% in fungi. Interestingly, phylogenetically aware, weighted metrics for fungal berry communities were not significantly different between regions. This could indicate that community differences caused by abiotic factors are based on the presence/absence of distinct strains belonging to the same microbial clades.

**Figure 3.**
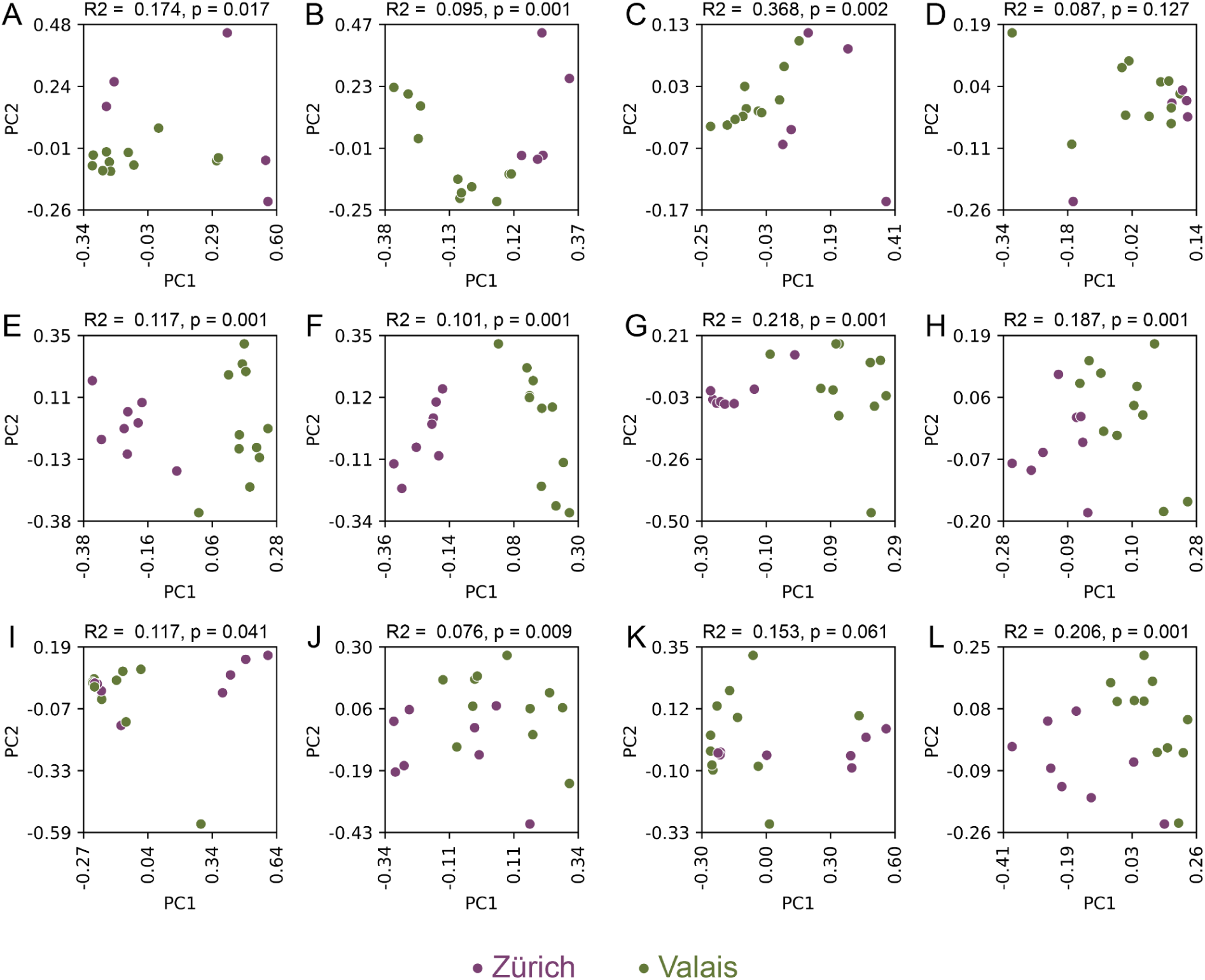
The two cantons (Valais in green, Zürich in purple) exhibit distinct microbial community composition as shown for soil (A-H) and berries (I-L), for bacterial communities (A-D) and fungal communities (E-L) with the beta diversity metrics of Bray Curtis (A, E, I) and Jaccard (B, F, J) and the k-mer based Bray Curits (C, G, K) and Jaccard (D, H, L). R2 and p-values are calculated with permutational analysis of variance (PERMANOVA) for the region.

Given the strong regional separation of microbial communities, we analyzed their correlations with abiotic factors separately, as well as within each region using a nested, multivariate permutational analysis of variance (PERMANOVA). Specifically, we assessed the association of soil microbiomes with soil properties (Supplementary Table 4), vineyard management practices (Supplementary Table 5) and site specific environmental factors (Supplementary Table 6). Fungal communities varied significantly with soil pH across all metrics. When nested for region, the clay content also significantly influenced fungal soil communities on lower taxonomic levels, indicating the subtle genetic or functional adaptation of fungi to specific soil textures (Supplementary Table 4). Bacterial communities varied strongly with pH, clay, and soil organic matter, and as a result, the overall microbial activity (CO_2_ respiration). These associations were substantially stronger when abundance was considered. Interestingly, the phylogenetically aware presence/absence metric (k-mer-based Jaccard) showed no significant differences by region in the multivariate PERMANOVA. This suggests that bacterial soil communities carry a robust phylogenetic structure and that the regional variation initially observed is largely mediated by soil properties that select for growth of particular clades; once these factors are accounted for, the regional effect disappears. These findings underscore the importance of including key soil variables when assessing drivers of microbial diversity in soil. Regarding the vineyard management practices we observed significant differences in the presence / absence of soil microorganisms with fertilizer and fungicide application (Supplementary Table 5). For bacterial communities, differences in abundance were also detected across different farming systems within a region. Among the site-specific environmental factors considered (altitude, annual rainfall, and average soil temperature), only the presence / absence of bacteria was significantly influenced by temperature and altitude (Supplementary Table 6). Notably, the relationship with altitude appears to be robust, as bacterial abundance also showed a significant association when analyzed within individual regions.

Similarly, in fungal communities of berries, we investigated associations with management practices (Supplementary Table 7) and environmental factors (Supplementary Table 8). Berry-associated fungi exhibit a significant association between fertilizer application and the presence/absence of strains (Supplementary Table 7). Of the site-specific factors, temperature significantly influenced the abundance and phylogenetic diversity of fungi when nested within the region. This is interesting as it indicates a robust, yet region-dependent effect of temperature on fungal berry communities.

### Soil Enzymatic Activity Reflects Microbial Function and Regionally Variable Soil Properties

Soil enzymatic activity varies with climatic conditions, soil properties and vineyard management and plays a major role in nitrogen availability and ultimately grape quality (Pawlowski *et al*. 2024). We analysed five key soil hydrolytic enzymes which are involved in the degradation of organic matter: β-Glucosidase (GLS), which catalyzes the hydrolysis of cellulose; β-Xylosidase (MUX), which hydrolyzes hemicellulose; β-N-Acetylglucosaminidase (NAG), which hydrolyzes chitin; Leucine aminopeptidase (LAP), which hydrolyzes polypeptides; and Acid/alkaline phosphomonoesterase (MUP), which hydrolyzes phosphate monoester bonds (Fetzer *et al*. 2025). All soil enzymatic activities, except MUX, varied significantly between the regions (see Supplementary Figure 2). Enzymatic activities were strongly and positively correlated with one another, apart from LAP which is negatively, but not significantly correlated with the other four (Figure 4A). CO_2_ emissions, which are a proxy for microbial activity, and enzymatic profiles were closely associated with key soil characteristics, many of which – particularly clay and total nitrogen content – vary significantly between regions. For instance, soil organic matter, which serves as a substrate for microbial growth, was expectedly strongly correlated with increased soil enzymatic activity. Similarly, clay content showed a strong association, as clay can also stabilize enzymes, protecting them from degradation and increasing their persistence. While increased CO_2_ emissions are associated with an increase across all measured enzymatic activities, increased microbial biomass – as determined by microbially bound nitrogen – was primarily associated with increased enzymatic activity regarding the degradation of cellulose and hemicellulose.

**Figure 4.**
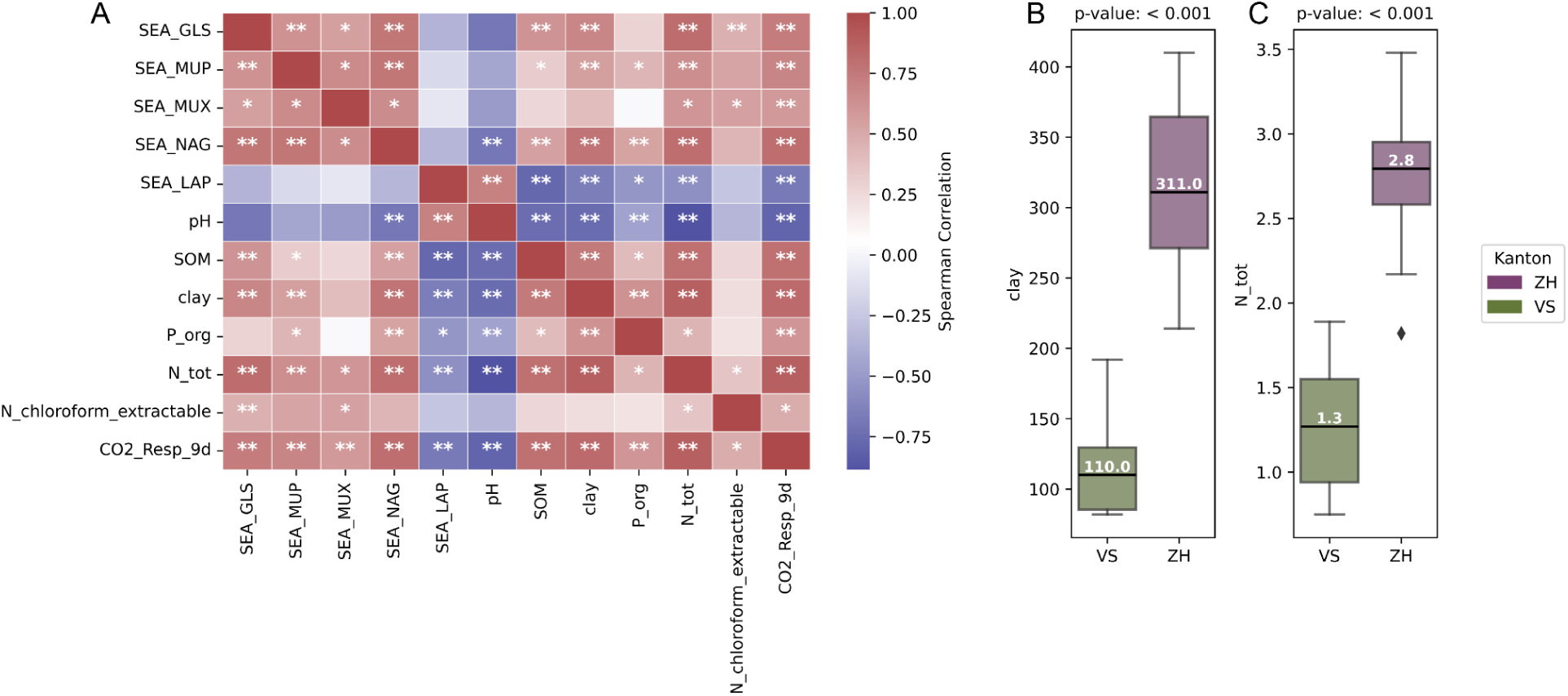
A. Spearman correlation heatmap of soil enzyme enzymatic activities (SEA), namely β-Glucosidase (GLS), Acid/alkaline phosphomonoesterase (MUP), β-Xylosidase (MUX), β-N-Acetylglucosaminidase (NAG), and Leucine aminopeptidase (LAP), and key soil properties (pH, soil organic matter, clay, organic phosphorus, total nitrogen, microbially bound chloroform-extractable nitrogen, CO2 respiration rate). (B) Clay and (C) total nitrogen (in g/kg dry matter of soil) variation between the regions (VS: Valais, ZH: Zürich).

### Soil pH and Microbial Activity are Key Distinguishing Soil Characteristics

We used Multiple Factor Analysis (MFA) (Escofier and Pagès 1994) to disentangle the influence and interactions of various factors in soil systems. The MFA integrates bacterial and fungal community profiles, climate (annual rainfall and average soil temperature), management practices, plant nutrient concentrations in leaves, altitude, soil enzymatic activities, and soil properties. The model captured 59.48% of total variance across seven components, indicating substantial explanatory power. Regional differences are the dominant driver distinguishing samples across all datalayers (see Figure 5). The first dimension (explaining 18.44% of variance) is primarily driven by soil pH and microbial activity, which display an inverse relationship. Additionally, microbially bound phosphorus and the relative abundance of a bacterial OTU from the uncultured genus of *Rokubacteriales* contributed significantly to this axis. The second dimension (10.28% of variance) was largely associated with altitude. When evaluating the absolute contributions of each group of variables, bacterial and fungal communities contributed the most to the overall model, likely due to their high feature counts and site-specific heterogeneity. Notably, their influence spanned across all dimensions of the MFA, reflecting the complexity of their interactions and connections with all other data groups in the model. Interestingly, altitude, soil enzymatic activities, and soil physicochemical properties contributed most prominently to the first dimension, reinforcing their roles in shaping the primary environmental gradient. In contrast, vineyard management exerted a strong influence on the second dimension, and was positioned farthest from the other groups in the MFA space (group representation), indicating a distinct effect that may act independently or modulate other variable groups. Among the management variables, disease management – specifically the use of synthetic fungicides – had the strongest influence, followed by fertilizer application and the cropping system (organic vs. conventional).

**Figure 5.**
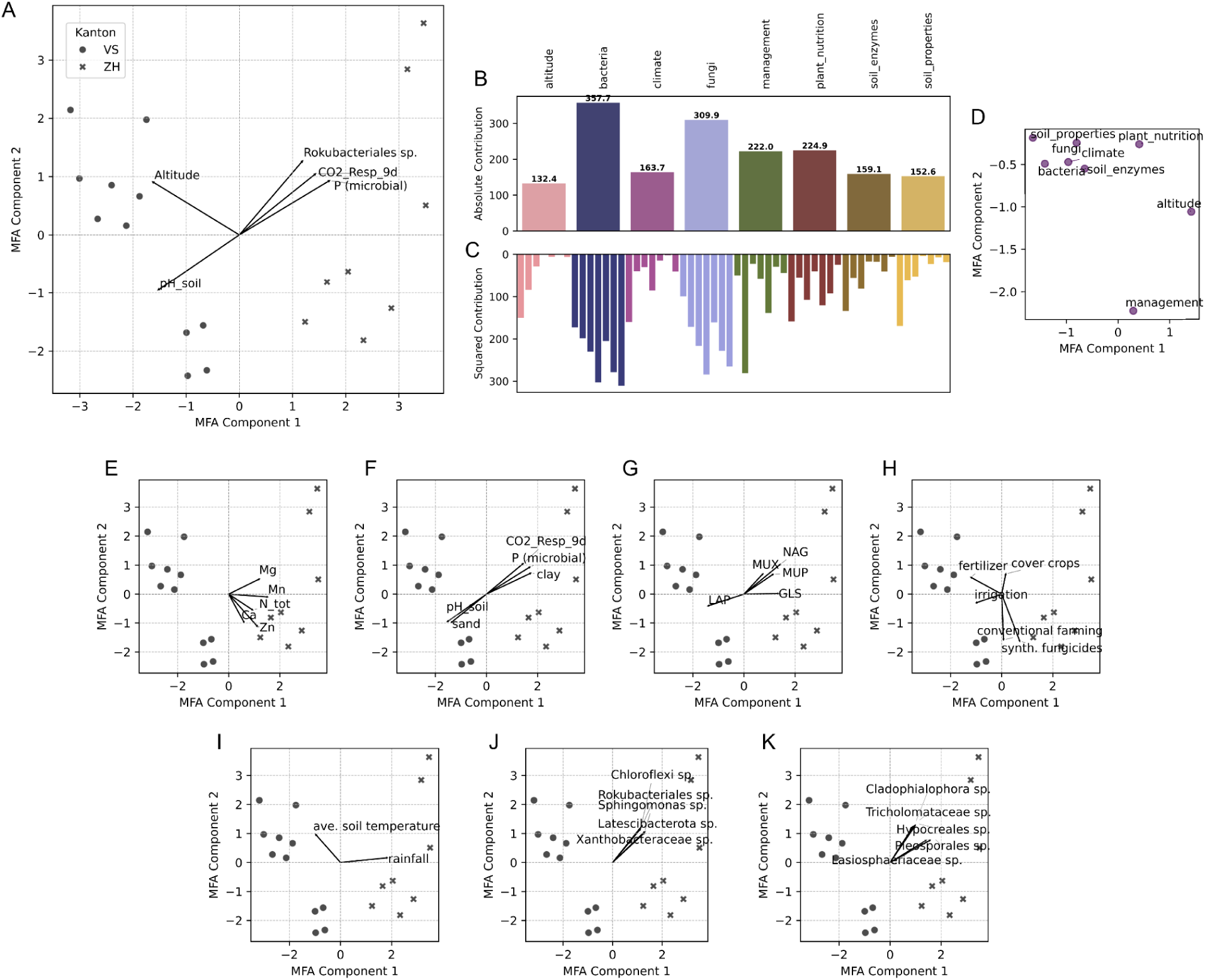
Multifactor analysis (MFA) of soil data layers showing the biplot with top 5 loadings (A), the absolute contributions to the model per group (B) and per dimension (C), as well as group representation per component (D). Separate biplots (E-I) show the top 5 loadings per group, namely within (E) plant nutrition measures, (F) soil properties, (G) soil enzymes, (H) management practices (I) climate, (J) bacteria and (K) fungi.

**Figure 6.**
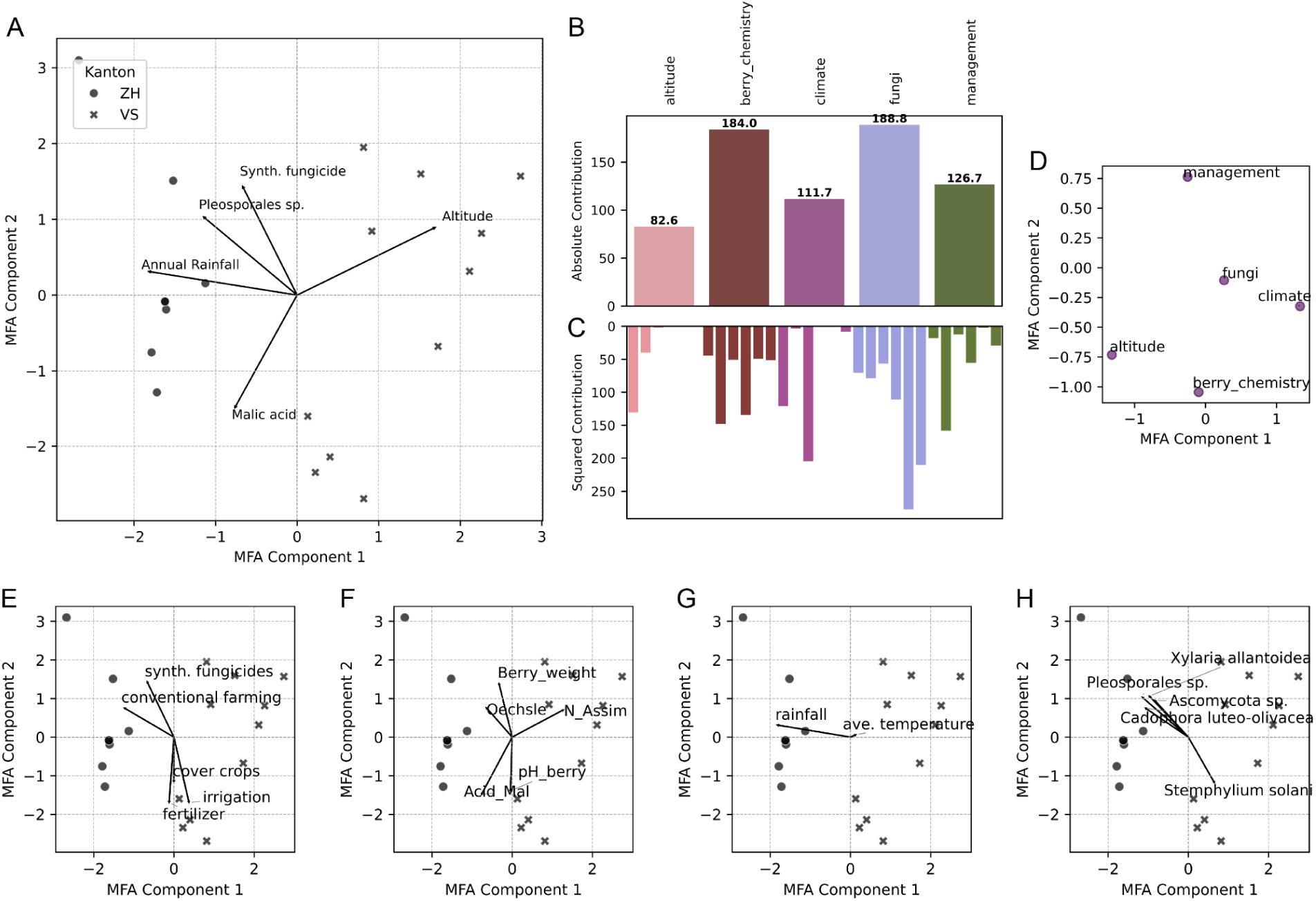
Multifactor analysis (MFA) of berry datalayers showing the biplot with top 5 loadings (A), the absolute contributions to the model per group (B) and per dimension (C), as well as group representation per component (D). Separate biplots (E-H) show the top 5 loadings per group, namely within (E) management practices, (F) berry chemistry, (G) climate, and (H) fungi.

### Integrative Analysis Highlights the Influence of Region, Altitude and Management on Berry Traits

We applied MFA to examine the relationships between berry chemistry, climate (annual rainfall, average air temperature), fungal communities, vineyard management practices, and plot altitude. The model, which used seven components, explained 70.87% of the variance, with 19.29% in dimension 1 and 17.47% in dimension 2. Similar to our observations for soil, region emerged as the primary distinguishing factor, with altitude being a dominant driver of the first dimension. The effect of altitude is mediated by a variety of factors, such as temperature or solar radiation, and we observe an inverse relationship with sugar content in berries (measured in degrees Oechsle). The contributions of climate and attitude to dimension 1 further underscore this relationship. Berry chemistry was a substantial contributing group to the overall variance in the model, following fungal communities, which exhibited the largest absolute contribution. This finding is noteworthy given that sugar content did not differ significantly between regions, as samples were collected at the same ripeness stage (see Supplementary Figure 3). The variability in sugar content, especially within canton Zürich (ranging from 75.1 to 103 °Oechsle), highlights intra-region variability at the same sampling date. For dimension 2, vineyard management emerged as the most important factor distinguishing samples. Specifically, fertilizer application and cover crop usage exhibited an inverse relationship in the MFA biplot, revealing potentially opposing effects. Interestingly, the group representation showed that all variables were distinctly separated in the MFA space, suggesting clear differentiation between factors and little overlapping structures in the individual datasets. Despite this separation, the model still explains more than 70% of overall variance, demonstrating the value of combining these data layers in comprehensively capturing the overall structure.

The large absolute group contribution, as well as the high contributions across dimensions of berry chemistry in the MFA, indicate that these measurements are influenced by various factors in the system. Interestingly, we did not observe direct, significant association of berry chemistry measures with individual site specific factors, such as altitude or climatic variations (Supplementary Figure 3). However, several noteworthy correlations with soil properties emerged. In particular, we noted a consistent pattern linking soil pH, microbial activity, organic phosphorus, and total nitrogen in the soil to tartaric acid concentration, likely causing the associated drop in pH and an increase in total acidity. Furthermore, soil organic phosphorus was negatively correlated with assimilable nitrogen in the berries. Correlation analysis further revealed that assimilable nitrogen content in berries is positively associated with both total nitrogen (Spearman’s ρ = 0.602, p = 0.0064) and calcium levels (ρ = 0.530, p = 0.0196) measured in the leaves. These associations reflect both the collinearity among factors and the interconnected roles of soil properties, plant nutrition and microbial function in shaping grape quality.

### Abundance and Presence of Fermentative Yeasts in Ripe Berries Correlates with Organic Acids

Berry chemistry parameters are key indicators of grape quality and play a critical role in shaping fermentation outcomes, ultimately influencing the characteristics of the resulting wine. To further investigate potential links between grape chemistry and fermentation, we analyzed the fungal microbiome of grape berries, which can also impact fermentation dynamics. Although the berry chemistry did not significantly influence the overall structure of the fungal community (Supplementary Table 9), we examined the presence and abundance of native fermentative yeast species of berries (*Hanseniaspora uvarum* and *Saccharomyces cerevisiae)*, whose presence/absence on berries did not differ significantly between regions (ANOVA: *H. uvarum*, BH-adjusted p = 0.118; *S. cerevisiae*, BH-adjusted p = 0.746). Interestingly, the abundance of these fermentative species was not significantly correlated with sugar content of the berries (Figure 7). However, we observed significant correlations (P < 0.05) of the two species with malic and tartaric acid concentrations, which are positively correlated with *H. uvarum* CLR-transformed abundance, and negatively with *S. cerevisiae* (Figure 7).

**Figure 7.**
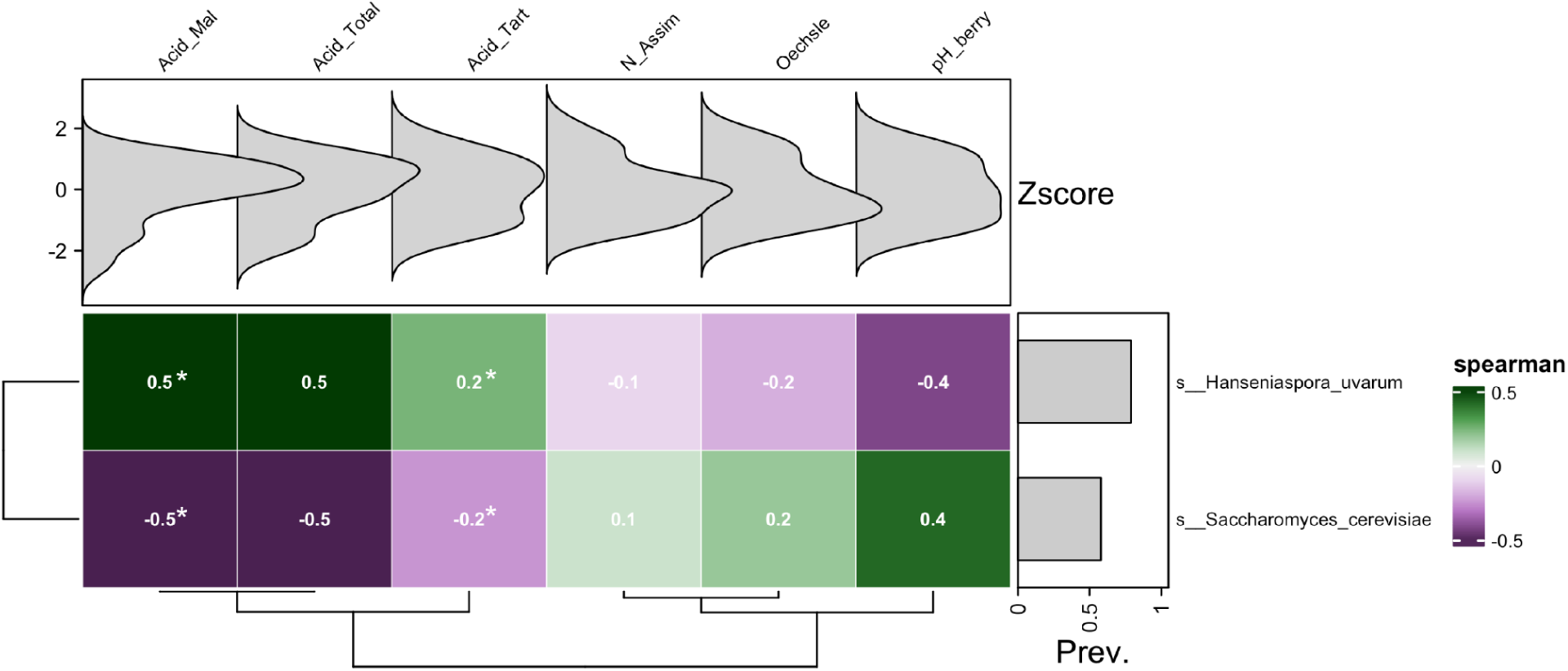
Testing the correlation of fermentative yeast species Hanseniaspora uvarum and Saccharomyces cerevisiae (CLR transformed abundance) with the various berry chemistry parameters (malic acid, total acidity, tartaric acid, assimilable nitrogen, sugar content in degree Oechsle and pH). (Asterisks denote p-values < 0.05 and are considered significant)

## Discussion

Viticulture is deeply rooted in tradition and local practices, therefore, a participatory citizen science approach can be valuable to integrate the objectives of winegrowers and researchers aiming to study agroecosystems. Involving local stakeholders can make sample collection more cost- and resource-effective, and enrich the scientific process through local knowledge and community engagement. In our project, workshops and active collaboration with winegrowers highlighted practical priorities, such as developing more site-specific solutions to support sustainable viticulture. This inclusive approach also revealed regional differences in viticultural practices which are shaped by tradition, geography, and local environment. Most winegrowers involved in this study produce wine from their vineyards themselves and showed a strong interest in further understanding the connections between vineyard health, soil conditions and resulting wine qualities. As research progresses towards developing more sustainable viticultural practices and understanding their effectiveness in a given environment, it is equally important to consider winegrowers’ practical needs and motivations for adopting new practices. Our study shows that, for instance, anticipated consumer appreciation and site-specific suitability, especially in the context of climate change, are key factors influencing the choice to plant new grape varieties. In contrast, financial incentives through subsidies, such as those introduced in Switzerland in 2022 to plant more resistant varieties (Der Schweizerische Bundesrat 2022), played a minor role.

Viticulture is arguably a particularly complex agricultural system – with grapes being highly sensitive to any environmental spatiotemporal variation, the high quality demands, as well as the complex long-term, labor intensive management and the integration of economic and cultural aspects. Any holistic research in this space therefore needs to be highly interdisciplinary and integrative of various datalayers (Giffard *et al*. 2022; Philippot *et al*. 2024). While some aspects of the viticultural system, such as the relationship between soil properties and wine quality, have been extensively studied, the intricate interactions between environmental and management factors remain less understood (Giffard *et al*. 2022). This is especially true when examining microbial diversity in vineyards. Microbial communities play a central role in viticulture and winemaking, as they contribute to distinct wine characteristics (Bokulich *et al*. 2014; Flörl *et al*. 2025c) and are shaped by a range of abiotic and biotic factors (Griggs *et al*. 2021), including the vineyard management (Burns *et al*. 2016; Morrison-Whittle, Lee and Goddard 2017; Chou *et al*. 2018; Coller *et al*. 2019; Vitulo *et al*. 2019). The efficacy of these management practices must be considered in the context of the cultivar (Steiner, Grace and Bacher 2021) and environment (Winter *et al*. 2018; Steiner *et al*. 2023). For example, in dry regions, traditional practices aim to suppress weeds in vineyards, through tillage or herbicides, to minimize water competition with the vines (Giffard *et al*. 2022). Conversely, in regions with higher precipitation and more nutrient-rich soils, preserving cover crops can be beneficial, as it can help prevent erosion and nutrient leaching during intense rainfalls (Giffard *et al*. 2022). Similarly, studies from different environments contrast in their effect of management on soil enzymatic activities (Peregrina, Pilar Pérez-Álvarez and García-Escudero 2014; Blondel *et al*. 2025). Our study, therefore, compared vineyards planted with the same cultivar, but situated in two climatically and pedologically distinct regions where we explored microbial diversity and function in soil and berries.

Despite differences in vineyard management, regional variations in altitude and climate remained the primary factor driving variation across all data layers, including the bacterial and fungal communities of soils and berries, nutrients concentration in leaves, soil enzymatic activities, and soil properties. While many of these measurements exhibited correlations with each other, our integrative approach revealed the central roles of pH and CO_2_ emissions from the soil, as well as a secondary strong effect of management, specifically the application of fungicides. In contrast, the analysis of berry samples, including berry chemistry and fungal communities, revealed that cover crop usage and fertilizer application were among the main drivers. Importantly, these distinct management effects were only detectable through an integrative analysis combining all data layers, as separate datasets often did not reveal these associations. This highlights the complexity of microbial systems and the interplay of factors shaping them. This complexity may also explain the mixed results of studies examining the link between vineyard microbiomes and management practices, such as cover crop usage (Novara *et al*. 2020; Wright *et al*. 2022; Li *et al*. 2023; Teixeira *et al*. 2025), cropping systems (Burns *et al*. 2016; Vitulo *et al*. 2019), fertilizer (Chou *et al*. 2018), tillage (Burns *et al*. 2016; Pingel, Reineke and Leyer 2019), or pesticide application (Agarbati *et al*. 2019; Steiner *et al*. 2024). While findings vary, for example whether cover crops influence soil or berry microbiome richness (Chou *et al*. 2018; Li *et al*. 2023; Teixeira *et al*. 2025), our analysis aligns with the broader conclusion that environmental factors are the primary determinants, with management practices acting as a secondary, yet distinct, influence in shaping microbial communities in soil and fruit.

To explore the effects of the regional variations on grape quality and potentially wine characteristics, we further examined key berry chemistry parameters and associated fungal communities. While direct correlations between individual site-specific factors and berry chemistry parameters did not show any significant associations, the multivariate model identified altitude and sugar content as primary drivers of variation in berry traits. This supports the notion that these traits are not shaped by isolated variables alone, but rather shaped by the combined effects of environmental and management factors. Additionally, we demonstrated a direct connection between soil properties, like the soil pH and CO_2_ emissions , with resulting berry chemistry, highlighting the role of the soil microbiome in influencing grape quality. We also identified associations between key fermentative yeast species and varying berry chemistry parameters, which is likely also connected to variable ripening dynamics between locations. Notably, *Saccharomyces cerevisiae* and *Hanseniaspora uvarum* exhibited a negative correlation with one another, suggesting niche competition. While similar patterns have been reported previously (Huang *et al*. 2022; Flörl *et al*. 2025c), the causal link between these dynamics and organic acid content in berries remains unclear and may reflect differences in metabolic capabilities or ecological niches between the species, or could result from broader community shifts during berry ripening.

A key limitation of this study is its cross-sectional design, offering only a snapshot of highly dynamic processes. Longitudinal sampling along and over multiple growing seasons will be critical to disentangle temporal variation and better understand cause-effect relationships. While we measured five key enzymatic activities, they only represent a fraction of the functional complexity in soil, and are solely a proxy for the microbial potential to hydrolyze organic compounds. For example, while we measured nitrogen levels in both soil and leaves, the mechanisms and timing of nitrogen availability in this context remains unresolved. Soil nitrogen must first be mineralized by microbial extracellular enzymes to become plant-accessible, yet we cannot determine from our data whether the observed nitrogen pools or enzymatic activity are recent or long-standing, nor how they fluctuate throughout the season.

Additionally, management practices in our study were assessed at a relatively broad level. For example, while we distinguished between cover crops and bare soil, we did not differentiate between underlying practices such as tillage versus herbicide use, or between spontaneous and sown cover crops. The complexity of vineyard management, including mixed approaches and influences by climate, timing of interventions, and vine age (Reynolds 2022), makes its specific influence particularly challenging to study. Furthermore, practices known to impact berry quality, such as canopy trimming or leaf removal (Spring *et al*. 2012; Reynolds 2022), were not assessed and would require larger, controlled experiments. Finally, the number of sites per region in our study was limited. Future work should expand sampling across more vineyards under clearly defined and comparable conditions to improve statistical power and capture regional heterogeneity more effectively.

In summary, our study clearly demonstrates the strong environmental dependency and interconnectedness of factors shaping microbial communities in vineyards, and their functional roles in vine nutrition and grape quality. These findings underscore the importance of integrating diverse datasets to understand such complex agroecological systems. By assembling multiple layers of environmental, biological, and management-related data, we contribute valuable pieces to a much larger and more intricate puzzle – how genotype, environment (including complex interactions between the microbiomes and plant host), and management interact to shape wine quality traits. These insights have the potential to guide the development and transition towards more sustainable and microbiome-informed vineyard management practices in the future.

## Methods

### Stakeholder Meetings, Questionnaire Development and Dissemination

First, we held co-design workshops in Zürich and Pully with a representative stakeholder group including winegrowers practicing different management approaches (biodynamic, conventional, organic) and other stakeholders from industry and government (e.g., from the organization of organic agriculture in Switzerland, BioSuisse, and the Swiss federal agriculture research organisation, Agroscope). These workshops served multiple functions: (i) to evaluate which research questions are of most interest to various stakeholder groups and (ii) to assess feasibility of a citizen science project (e.g., winegrower willingness to answer a questionnaire or perform sampling) as well as (iii) to establish networks and contacts of winegrowers interested in participating in the project. The workshops were held in the French- and German-speaking parts of Switzerland, with 20 and 21 participants, respectively. Winegrowers identified shared key research priorities, such as the role of the soil and grape microbiome, practical alternatives to chemical inputs, and the economic and ecological impacts of sustainable practices. Participants emphasized the need for clearer definitions of sustainability, more site-specific solutions, and stronger exchange between researchers and practitioners to support informed decision-making in the vineyard. Based on the consensus between winegrowers’ practical priorities and current frontiers of microbial ecology research, we launched a project focusing on the impact of management effects on soil and grape microbiota.

With the input generated in the co-design workshops we developed a questionnaire, both in German and French (available at Zenodo: 10.5281/zenodo.16418780). The questionnaire was distributed through the established network and received 80 responses, 35 in German and 45 in French, of which 6 were incomplete.

We surveyed viticultural practices and analysed these by cultural-linguistic region, as vineyard management is influenced by climatic and geographical conditions, as well as local traditions. Respondents grow between 1 and 33 different varieties each, with an average of 5.1 red and 5.3 white varieties per winegrower (Supplementary Figure 1B). The most popular red grape varieties were similar across regions (Pinot Noir and Merlot), but preferences for white varieties differed. In French-speaking regions, Chasselas, Savagnin blanc and Petite Arvine were the most popular, while winegrowers from German-speaking regions preferred Müller-Thurgau, Chardonnay and Sauvignon Blanc (Supplementary Figure 1C-D).

Analysis of the questionnaire was performed in R (version 4.5.0), using the package tidyverse (Wickham *et al*. 2019).

### Vineyard Sites and Sample Collection

A total of 19 guyot-trained Pinot Noir vineyards in Switzerland were analysed, comprising 11 sites in the Rhône Valley of Canton Valais (Calcisol soil) and 8 sites in Canton Zürich (Cambisol soil). Four field trips were conducted: in Valais at veraison on August 17 and at harvest on August 30, 2023; and in Zürich on September 5 and 14, 2023 respectively. At veraison we sampled soil and leaves, and collected ripe berries at harvest. Soil was collected from three locations along a single row, at a depth of 0–30 cm. Soil samples were sieved (2 mm) and pooled per plot and kept moist at 5°C. 25 fully developed leaves were collected along the same row for nutrient analysis. At harvest, 300 grape berries were randomly collected from different grape clusters within the same vineyard rows. For microbiome analysis grape samples were obtained by excising berry clusters with sterile scissors. All samples were transported on ice to the laboratory and stored at −80 °C until further processing.

### Climate Data

Meteorological data, including annual precipitation, mean annual air temperature at 2 m above ground level, and mean annual soil temperature at 10 cm depth, were obtained from weather stations in the Agrometeo database (Agroscope). Data were collected for the period from September 1, 2022, to August 31, 2023. For each vineyard, the nearest available weather station was selected, with an average distance of 1.8 km between the vineyards and the corresponding stations.

### DNA extraction, library preparation, sequencing

Samples were processed as previously described (Flörl *et al*. 2025c). Briefly, DNA was extracted using the MagAttract PowerSoil Pro DNA Kit (QIAGEN, cat. no. 47109) and the PowerBead Pro Plate (QIAGEN, cat. no. 19311). For library preparation we used the HighALPS ultra-high-throughput library preparation protocol (Flörl *et al*. 2024). Bacterial communities were profiled by sequencing the hypervariable V4 region of the 16S rRNA (primers: 515F (Parada, Needham and Fuhrman 2016) and 806R (Apprill *et al*. 2015)). For fungal community analysis, the first locus of the internal transcribed spacer (ITS) region was amplified using the BITS and B58S3 primers (Bokulich and Mills 2013). To reduce host plastid and mitochondrial 16S rRNA gene contamination we used peptide nucleic acid (PNA) clamps (Lundberg *et al*. 2013; Flörl and Bokulich 2024). The libraries were sequenced at the Functional Genomics Center Zürich on a Illumina NextSeq 2000 P1 (600 cycles) with 20% PhiX. In total, 1’361’332 reads were obtained.

### Microbiome Data Analysis

Microbial diversity analysis was conducted using QIIME 2 version 2024.10 (Bolyen *et al*. 2019). Demultiplexed Illumina fastq files were imported, and any residual adapters trimmed using cutadapt (Martin 2011), followed by denoising with DADA2 (Callahan *et al*. 2016). For 16S, paired-end reads were processed, while single-end reads were used for ITS. The resulting amplicon sequencing variants (ASVs) were taxonomically assigned using the q2-feature-classifier plugin (Bokulich *et al*. 2018), employing a naïve Bayes taxonomy classifier trained on the 99 % SILVA (Quast *et al*. 2012) 16S rRNA gene database (138 release), trimmed to the 515F-806R (V4) region (Robeson *et al*. 2021) and weighted to better represent plant-surface communities (Kaehler *et al*. 2019). Similarly, fungal ASVs were classified using the UNITE database (v9.0, Version 18.07.2023) (Abarenkov *et al*. 2023). Contaminants were identified and removed with decontam (Davis *et al*. 2017), using DNA extraction negative controls. Additionally, non-target ASVs were filtered out, including fruiting body-forming mushrooms from ITS and host DNA (mitochondria, chloroplasts) from 16S, along with ASVs that could not be classified at least to the phylum level. To assess microbial diversity, samples were rarefied for bacteria to 500 reads in soil samples and in fungi to 5’000 reads for soil samples and 10’000 reads in berry samples. Alpha diversity (Pielou evenness (Pielou 1966), observed features, Shannon entropy (Shannon 1948)) and beta diversity metrics (Bray Curtis (Sorensen Thorvald 1948), Jaccard (Jaccard 1908)) were computed using the q2-diversity plugin. In parallel, diversity estimates were also derived from k-mer profiles (Bokulich 2025). To evaluate differences in community composition, permutational multivariate analysis of variance (PERMANOVA) was carried out via the ADONIS function (Anderson 2001), implemented in q2-diversity. Multifactor analysis (MFA) was done using the prince package (Halford 2025) and therefore we clustered ASVs against the reference databases at 90% identity for fungi and 99% for bacteria. Visualizations were produced with microViz (Barnett, Arts and Penders 2021), matplotlib (The Matplotlib Development Team 2024), and seaborn (Waskom 2021), while geographic mapping was done with geopandas (Jordahl *et al*. 2020).

### Leave Plant Nutrition Analysis

Leaf nutrient content was analyzed by Sol-Conseil (Gland, Switzerland) for nitrogen (N), phosphorus (P), potassium (K), calcium (Ca), magnesium (Mg), copper (Cu), iron (Fe), manganese (Mn), zinc (Zn), and boron (B). The methods used by this laboratory are described under https://sol-conseil.ch/extraits-de-methodes/.

### Soil Properties

Soil pH (H₂O method), soil organic matter (SOM), soil texture, CaCO₃ content, and water-soluble concentrations of phosphorus (P), potassium (K), calcium (Ca), and magnesium (Mg) (H₂O10 method) were analyzed on 2 mm sieved air dried samples by Sol-Conseil (Gland, Switzerland), a certified and SAS-accredited soil laboratory. The methods used to acquire these data are further described on the Sol-Conseil website (https://sol-conseil.ch/extraits-de-methodes/). Additionally, Total carbon (TC), total nitrogen (TN), organic phosphorus (Porg), and maximum water-holding capacity (WHC) were measured in the group of plant nutrition at ETH Zurich. To minimize analytical variation, dried soil samples were finely ground before TC and TN content was determined via combustion analysis (FlashEA Analyzer, Thermo Fisher Scientific). Porg was quantified using the Saunders and Williams method (Saunders and Williams 1955). In brief, 2 g of air-dried soil sieved at 2 mm was combusted in a porcelain crucible at 550 °C for one hour. Both ashed and unashed samples (2 g each) were extracted with 25 ml of 0.5 M H₂SO₄ for 16 hours on a horizontal shaker. The extracts were filtered through 0.2 μm Millipore filters, and reactive phosphate concentrations were determined colorimetrically at 610 nm using the malachite green method (Ohno and Zibilske 1991). The increase in H₂SO₄-extractable phosphorus following ashing was attributed to organic phosphorus (Porg). Water-holding capacity (WHC) was determined also on 2 mm sieved soil according to the B-WHK protocol (Agroscope 2020).

### Soil Microbiological Activity

Prior to incubation, fresh soil samples were brought at 60% of their maximum water-holding capacity (WHC) and 20°C in the dark for a week. The pre-incubated soils were then incubated in darkness at 20 °C (± 1 °C) for nine days.

Soil respiration (mg CO₂/kg dry soil) was measured at day 3, 6, and 9 of incubation following the method of Alef and Nannipieri (Alef and Nannipieri 1995). The CO₂ released from the soil was captured in NaOH and quantified via HCl titration.

Mineral nitrogen (NH₄⁺ and NO₃⁻) was analyzed at day 9 using the Nmin method by Sol-Conseil (Gland, Switzerland; https://sol-conseil.ch/extraits-de-methodes/). Hexanol-extractable phosphorus (P) in microbial biomass was determined at day 9 on moist soils using the hexanol fumigation-extraction method in combination with anion exchange membranes (Kouno, Tuchiya and Ando 1995). Chloroform extractable nitrogen (N) was estimated using the fumigation-extraction method (Vance, Brookes and Jenkinson 1987).

### Soil Enzymatic Activity

Soil enzyme activity was analyzed by DigitSoil (https://www.digit-soil.com/) on their Soil Enzymatic Activity Reader (SEAR) device which enables the simultaneous measurement of potential extracellular soil enzyme activity in fresh soil samples using a contact-based approach. Extracellular enzymatic activities of β-glucosidase, β-xylosidase, β-N-acetylglucosaminidase, leucine aminopeptidase, and acid/alkaline phosphomonoesterase were measured in approximately 15 g of incubated moist soil samples. The assay employed enzyme-specific fluorogenic substrates, including 4-methylumbelliferyl N-acetyl-β-D-glucosaminide, 4-methylumbelliferyl-β-D-glucopyranoside, 4-methylumbelliferyl phosphate, 4-methylumbelliferyl-β-D-xylopyranoside, and L-leucine-7-amido-4-methylcoumarin hydrochloride. Enzyme activity was quantified based on the increase in fluorescence over time, with each membrane containing methylumbelliferyl (MUF) and amido-4-methylcoumarin (AMC) standards for calibration. The method used by DigitSoil to measure extracellular hydrolytic enzymatic activities has been described in (Monteux *et al*. 2024) and (Fetzer *et al*. 2025).

### Berry Chemistry

The grape must was analyzed by Agroscope using infrared spectroscopy with the FOSS Winescan™ (Foss Electric, Hillerød, Denmark) and included measurements of sugar concentration (degrees Oechsle), pH, yeast-available nitrogen (YAN), total acidity as well as malic and tartaric acid concentration.

## Data Availability

Amplicon sequencing data is available from the Sequence Read Archive (SRA) under accession number PRJEB89111 (16S) and PRJEB89112 (ITS).

## Code Availability

All code used in this study is available on GitHub: https://github.com/LenaFloerl/WINE-CitizenScience

## Funding

This work was supported by a Citizen Science Zurich Seed Grant, from the Swiss National Science Foundation [Grant Number: 310030_204275] (to NAB), and the Swiss Government Excellence Ph.D. Scholarship (to LF).

## Acknowledgements

We would like to thank Citizen Science Zürich (ETH Zurich / University of Zurich) for their advice in implementing a co-design approach, Prof. Dr. Michael Siegrist for the valuable insights in questionnaire design, Fabian Regel who provided technical support in soil analyses, and the many winegrowers and stakeholders who participated in the workshops, answered the questionnaire, gave us feedback, and let us sample their vineyards.

## Conflicts of interest

None declared.

## Author Contributions

The study was conceived by NAB, LF, and EF, and supervised by NAB. LF, MPRM, JSR, VZ and KMH collected samples. VZ and KHM performed the berry chemistry analysis, the laboratory of EF measured soil biological activities and LF did the microbiome sequencing. LF performed the data analysis and MPRM evaluated the questionnaires. LF wrote the article with input from MPRM and NAB, and KMH, EF, and VZ reviewed and edited the article.

## Glossary

GLS: β-Glucosidase
MUX: -Xylosidase
NAG: β-N-Acetylglucosaminidase
LAP: Leucine aminopeptidase
MUP: Acid/alkaline phosphomonoesterase
MFA: Multifactor Analysis
Porg: Organic phosphorus
ASV: Amplicon sequencing variant
PERMANOVA: permutational multivariate analysis of variance

## Supplementary Information

- Supplementary Figures
- Supplementary Tables
- Questionnaires (English, German, French): at Zenodo under 10.5281/zenodo.16418780

## Notes

### Competing Interest Statement

The authors have declared no competing interest.

