## Supplementary Figures and Tables for "Interaction of Environment and Vineyard Management Effects Shape Microbial Functionality, *Terroir*, and Grape Qualities"

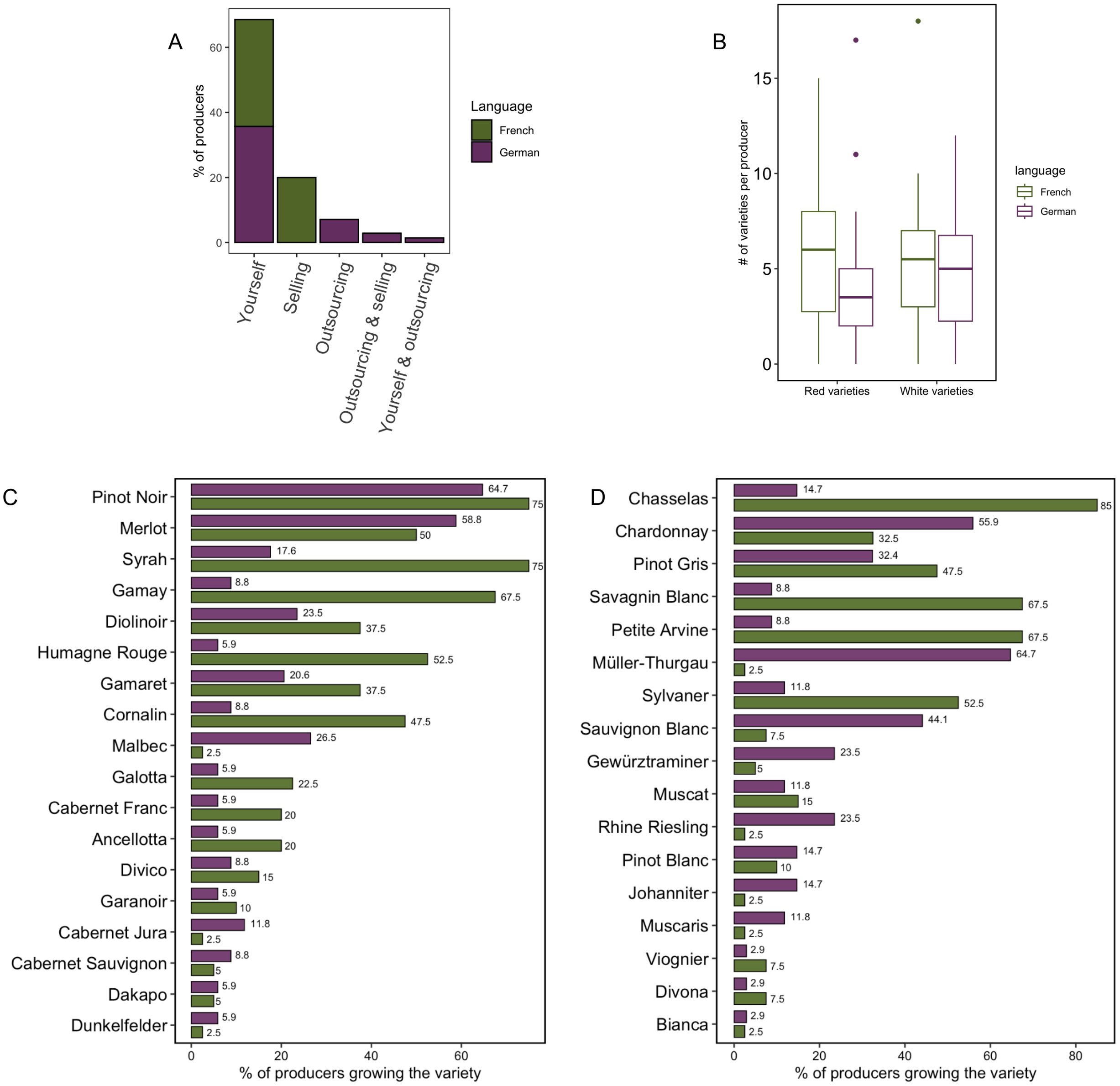


***Supplementary Figure 1.*** *(A) A majority of respondents produce their own wine, while smaller percentages sell their grapes, mostly in French speaking regions, or outsource the production.(B) The number of varieties grown by each producer is highly variable, but does not differ greatly between the two cultural-linguistic regions. (C, D) The most common white grape varieties differ between regions, while the most popular red varieties are the same across respondents from French and German- speaking Switzerland.*

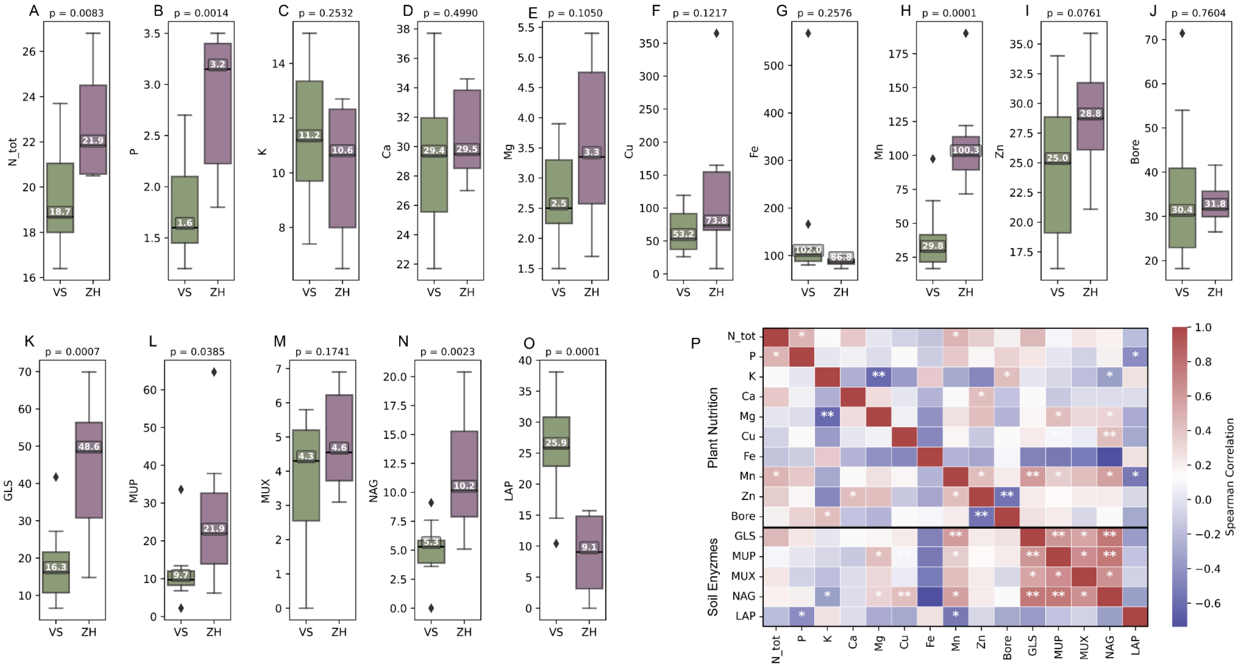


***Supplementary Figure 2**: Plant nutrients concentrations in vine leaves (A. total nitrogen, B. phosphorus. C. potassium, D. calcium, E. magnesium – all in g/kg dry matter; F. copper, G. iron, H. manganese, I. zinc, J. boron – all in mg/kg dry matter) as well as soil enzymatic activity in pmol/min (K. β-Glucosidase (GLS), L. Acid/alkaline phosphomonoesterase (MUP), M. β-Xylosidase (MUX), N. β-N-Acetylglucosaminidase (NAG) and O. Leucine aminopeptidase (LAP)) varying between the cantons Valais (VS) and Zurich (ZH), as well as the Spearman correlation between these factors. (p-values < 0.05 are considered significant and marked with one asterisks, p-values < 0.01 are marked with two asterisk)*

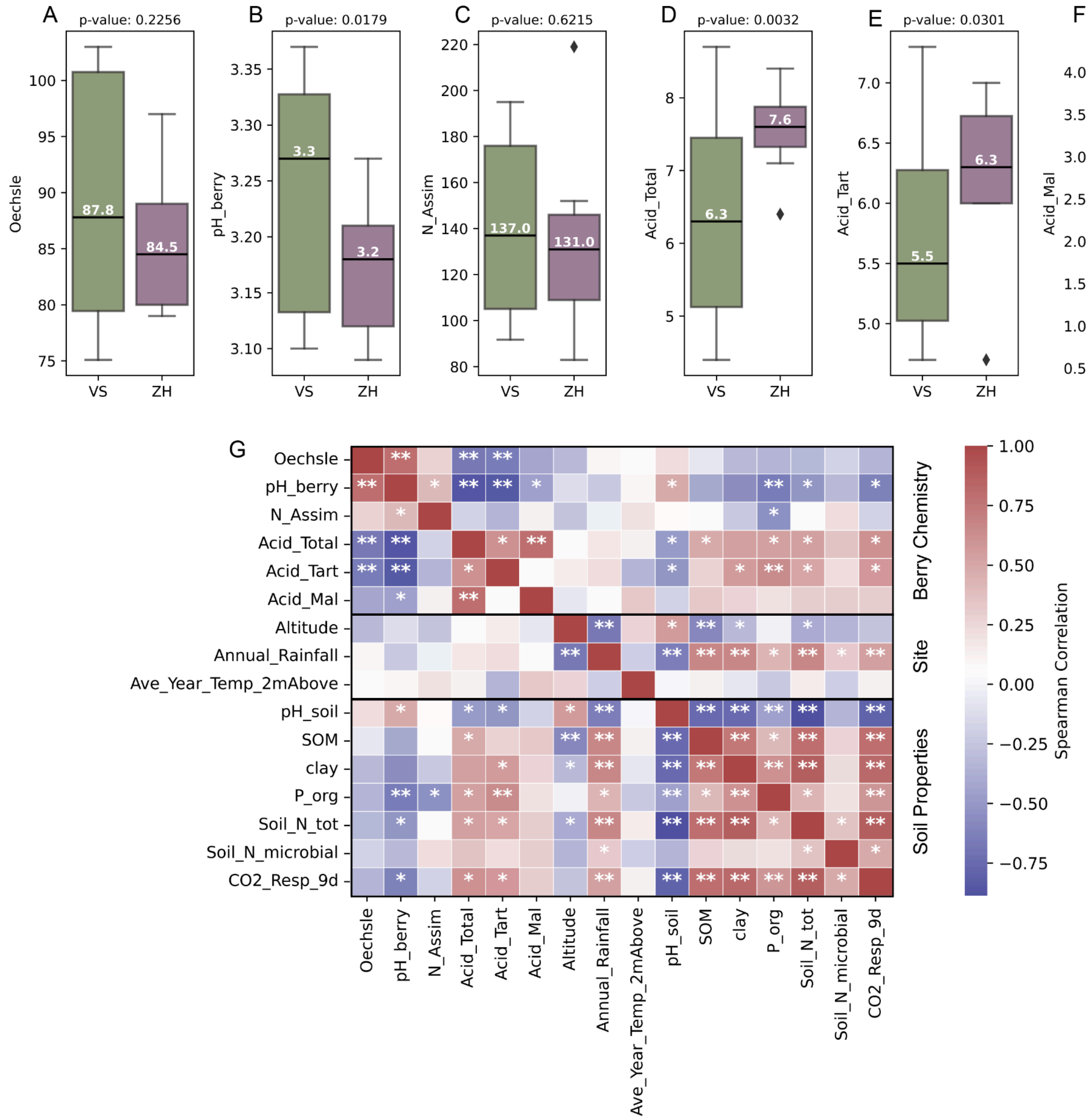


***Supplementary Figure 3****: A-F. Berry chemistry parameters varying between cantons Valais (VS) and Zurich (ZH) (specifically A. sugar content in degree Oechsle, B. pH of the berries, C. assimilable nitrogen in mg/L, D. total acidity in g/L, E. tartaric acid in g/L and F. malic acid in g/L). G. Spearman correlation between berry chemistry parameters and site specific environmental factors as well as various soil properties. (p-values < 0.05 are considered significant and marked with one asterisks, p-values < 0.01 are marked with two asterisk)*

### 1. Supplementary Tables

***Supplementary Table 1***: *Overview of the sampled vineyards in cantons Valais and Zurich and their respective management practices, soil properties, location and climate data.*

| Vineyard ID | Canton | Management | | | | | Soil Properties | | | | | | Location | | Climate | | |
|---|---|---|---|---|---|---|---|---|---|---|---|---|---|---|---|---|---|
| | | Cropping System | Fertilizer Usage | Irrigation | Soil Treatment | Disease Management | pH | Soil Organic Matter | Clay (%) | Silt (%) | Sand (%) | Max. Water Holding Capacity | Coordinates | Altitude (m) | Annual Rainfall (mm) | Ave. Temp 10cm Below (°C) | Ave. Year Temp 2m Above (°C) |
| 1_VS | Valais | conventional | no | no | bare soil | synthetic fungicides | 7.9 | 16 | 8.3 | 2.34 | 6.82 | 0.27 | 46.187, 7.222 | 510 | 716.7 | 12.742 | 12.067 |
| 2_VS | Valais | conventional | no | no | cover crops | synthetic fungicides | 7.8 | 19 | 8.5 | 2.05 | 7.11 | 0.32 | 46.187, 7.222 | 510 | 716.7 | 12.742 | 12.067 |
| 3_VS | Valais | conventional | no | yes | bare soil | synthetic fungicides | 8.1 | 24 | 8.2 | 2.65 | 6.54 | 0.27 | 46.187, 7.222 | 510 | 716.7 | 12.742 | 12.067 |
| 4_VS | Valais | conventional | no | yes | cover crops | synthetic fungicides | 8.1 | 25 | 8.6 | 2.64 | 6.50 | 0.26 | 46.187, 7.222 | 510 | 716.7 | 12.742 | 12.067 |
| 5_VS | Valais | organic | yes | no | cover crops | organic fungicides | 8.1 | 13 | 1.10 | 3.35 | 5.55 | 0.27 | 46.100, 7.058 | 710 | 939.9 | 12.708 | 12.075 |
| 6_VS | Valais | conventional | yes | yes | bare soil | synthetic fungicides | 7.9 | 19 | 1.38 | 3.83 | 4.80 | 0.33 | 46.243, 7.332 | 770 | 624.8 | 16.192 | 12.308 |
| 7_VS | Valais | organic | yes | yes | cover crops | organic fungicides | 7.9 | 20 | 1.21 | 3.33 | 5.45 | 0.42 | 46.244, 7.331 | 816 | 624.8 | 16.192 | 12.308 |
| 8_VS | Valais | conventional | yes | yes | bare soil | organic fungicides | 7.9 | 15 | 9.80 | 4.09 | 4.92 | 0.43 | 46.245, 7.374 | 731 | 729.8 | 13.692 | 11.158 |
| 9_VS | Valais | conventional | yes | no | cover crops | organic fungicides | 8.0 | 15 | 1.92 | 3.75 | 4.33 | 0.19 | 46.299, 7.524 | 695 | 703.5 | 13.908 | 11.817 |
| 10_VS | Valais | organic | yes | no | cover crops | organic fungicides | 7.8 | 33 | 1.13 | 3.19 | 5.68 | 0.42 | 46.309, 7.554 | 682 | 673.4 | 14.275 | 12.308 |
| 11_VS | Valais | conventional | no | yes | cover crops | organic fungicides | 8.1 | 20 | 1.45 | 3.78 | 4.77 | 0.43 | 46.316, 7.649 | 844 | 681.8 | 13.967 | 12.017 |
| 12_ZH | Zurich | conventional | yes | no | bare soil | synthetic fungicides | 7.5 | 32 | 2.14 | 3.40 | 4.46 | 0.50 | 47.570, 8.591 | 423 | 989.7 | 11.883 | 11.408 |

| | | | | | | | | | | | | | | | | |
|---|---|---|---|---|---|---|---|---|---|---|---|---|---|---|---|---|
| 13_ZH | Zurich | conventional | no | no | cover crops | synthetic fungicides | 7.8 | 42 | 2.92 | 3.99 | 3.09 | 0.57 | 47.514, 8.703 | 460 | 1160.8 | 12.242 | 11.367 |
| 14_ZH | Zurich | conventional | no | no | bare soil | synthetic fungicides | 7.6 | 55 | 3.63 | 3.40 | 2.98 | 0.67 | 47.247, 8.731 | 534 | 1274.3 | 12.85 | 12.108 |
| 15_ZH | Zurich | conventional | no | yes | cover crops | synthetic fungicides | 7.7 | 50 | 2.75 | 2.83 | 4.41 | 0.57 | 47.236, 8.749 | 421 | 1274.4 | 12.85 | 12.108 |
| 16_ZH | Zurich | organic | no | no | cover crops | organic fungicides | 6.9 | 57 | 4.10 | 3.47 | 2.43 | 0.75 | 47.252, 8.715 | 516 | 1274.3 | 12.85 | 12.108 |
| 17_ZH | Zurich | organic | no | no | bare soil | organic fungicides | 7.0 | 47 | 3.69 | 3.64 | 2.67 | 0.69 | 47.270, 8.674 | 475 | 1134.1 | 13.85 | 12.025 |
| 18_ZH | Zurich | conventional | no | no | cover crops | synthetic fungicides | 7.6 | 61 | 3.30 | 3.29 | 3.41 | 0.60 | 47.271, 8.637 | 422 | 1134.1 | 13.85 | 12.025 |
| 19_ZH | Zurich | conventional | no | no | bare soil | synthetic fungicides | 7.6 | 45 | 2.60 | 3.27 | 4.13 | 0.56 | 47.310, 8.594 | 473 | 1134.1 | 13.85 | 12.025 |

***Supplementary Table 2**: Alpha diversity metrics of fungal and bacteria communities in soil across the region and under different vineyard management strategies. (p-values < 0.05 are considered significant and highlighted in bold)*

| | Canton | Farming System: Organic / Conventional | Fertilizer application (yes / no) | Irrigation (yes / no) | Soil treatment (bare soil / cover crops) | Disease management (Fungicide application: yes / no) |
|---|---|---|---|---|---|---|
| | | | **ITS** | | | |
| **Pielou Evenness** | 0.457 | 0.355 | 0.353 | 0.933 | 0.430 | 0.620 |
| **Observed Features** | 0.869 | 0.405 | 0.447 | 0.800 | 0.079 | 0.069 |
| **Shannon Entropy** | 0.620 | 0.229 | 0.272 | 0.673 | 0.161 | 0.069 |
| | | | **16S** | | | |
| **Pielou Evenness** | **0.020** | 0.533 | 0.828 | 0.125 | 0.225 | 0.808 |
| **Observed Features** | 0.156 | 1.000 | 0.385 | 0.751 | 0.762 | 0.805 |
| **Shannon Entropy** | 0.062 | 0.955 | 0.448 | 0.315 | 0.332 | 0.893 |

***Supplementary Table 3**: Alpha diversity metrics of fungal communities of berries across the region and under different vineyard management strategies. (p-values < 0.05 are considered significant and highlighted in bold)*

| | Canton | Farming System: Organic / Conventional | Fertilizer application (yes / no) | Irrigation (yes / no) | Soil treatment (bare soil / cover crops) | Disease management (Fungicide application: yes / no) |
|---|---|---|---|---|---|---|
| **Pielou Evenness** | 0.741 | 0.643 | 0.673 | 0.499 | 0.930 | 0.804 |
| **Observed Features** | 0.321 | 0.228 | 0.352 | 0.236 | 0.219 | 0.679 |
| **Shannon Entropy** | 0.137 | 0.195 | 0.673 | 0.176 | 0.599 | 0.563 |

***Supplementary Table 4****: Nested, multivariate permutational analysis of variance (PERMANOVA) to assess correlations between fungal (ITS) and bacterial (16S) communities in soil with various soil properties, calculated per factor as well as nested by region. (p-values < 0.05 are considered significant and highlighted in bold)*

| | **ITS** | | | | | | | | **16S** | | | | | | | |
|---|---|---|---|---|---|---|---|---|---|---|---|---|---|---|---|---|
| | **Bray Curtis** | | **Jaccard** | | **Kmer: Bray Curtis** | | **Kmer: Jaccard** | | **Bray Curtis** | | **Jaccard** | | **Kmer: Bray Curtis** | | **Kmer: Jaccard** | |
| | **R2** | **p-val** | **R2** | **p-val** | **R2** | **p-val** | **R2** | **p-val** | **R2** | **p-val** | **R2** | **p-val** | **R2** | **p-val** | **R2** | **p-val** |
| **Canton** | 0.120 | **0.001** | 0.101 | **0.001** | 0.222 | **0.001** | 0.189 | **0.001** | 0.185 | **0.001** | 0.095 | **0.001** | 0.368 | **0.001** | 0.087 | 0.212 |
| **pH soil** | 0.064 | **0.025** | 0.061 | **0.028** | 0.079 | **0.040** | 0.090 | **0.018** | 0.179 | **0.001** | 0.075 | **0.012** | 0.084 | **0.028** | 0.066 | 0.515 |
| **clay** | 0.052 | 0.247 | 0.056 | 0.132 | 0.045 | 0.391 | 0.049 | 0.279 | 0.164 | **0.001** | 0.075 | **0.015** | 0.108 | **0.009** | 0.045 | 0.853 |
| **CO2 Resp.** | 0.055 | 0.134 | 0.056 | 0.114 | 0.053 | 0.223 | 0.042 | 0.385 | 0.057 | **0.033** | 0.063 | 0.260 | 0.039 | 0.119 | 0.101 | 0.157 |
| **WHC max** | 0.049 | 0.394 | 0.050 | 0.497 | 0.039 | 0.538 | 0.036 | 0.555 | 0.040 | 0.145 | 0.063 | 0.249 | 0.018 | 0.330 | 0.043 | 0.847 |
| **SOM** | 0.057 | 0.111 | 0.058 | 0.083 | 0.044 | 0.409 | 0.046 | 0.320 | 0.036 | 0.168 | 0.065 | 0.182 | 0.060 | **0.033** | 0.103 | 0.137 |
| **Canton : pH soil** | 0.043 | 0.693 | 0.048 | 0.713 | 0.052 | 0.240 | 0.047 | 0.275 | 0.057 | **0.044** | 0.067 | 0.120 | 0.083 | **0.019** | 0.070 | 0.417 |
| **Canton : clay** | 0.071 | **0.010** | 0.062 | **0.030** | 0.059 | 0.155 | 0.053 | 0.185 | 0.053 | 0.068 | 0.062 | 0.353 | 0.064 | 0.056 | 0.042 | 0.835 |
| **Canton : SOM** | 0.047 | 0.518 | 0.054 | 0.203 | 0.035 | 0.642 | 0.073 | 0.052 | 0.075 | **0.020** | 0.066 | 0.160 | 0.069 | **0.031** | 0.039 | 0.919 |
| **Canton : CO2 Resp** | 0.047 | 0.516 | 0.051 | 0.332 | 0.046 | 0.336 | 0.052 | 0.238 | 0.038 | 0.153 | 0.064 | 0.282 | 0.025 | 0.284 | 0.055 | 0.642 |
| **Canton : WHC max** | 0.058 | 0.095 | 0.051 | 0.378 | 0.032 | 0.709 | 0.041 | 0.441 | | | | | | | | |
| **Residuals** | 0.335 | | 0.353 | | 0.295 | | 0.282 | | 0.116 | | 0.303 | | 0.082 | | 0.350 | |

***Supplementary Table 5****: Nested, multivariate permutational analysis of variance (PERMANOVA) to assess correlations between fungal (ITS) and bacterial (16S) communities in soil with various vineyard management practices (Fertilizer application, Disease management: organic or synthetic fungicide application, Cropping System: organic vs conventional, Irrigation, Soil treatment: cover crops vs bare soil), calculated per factor as well as nested by region. (p-values < 0.05 are considered significant and highlighted in bold)*

| | **ITS** | | | | | | | | **16S** | | | | | | | |
|---|---|---|---|---|---|---|---|---|---|---|---|---|---|---|---|---|
| | **Bray Curtis** | | **Jaccard** | | **Kmer: Bray Curtis** | | **Kmer: Jaccard** | | **Bray Curtis** | | **Jaccard** | | **Kmer: Bray Curtis** | | **Kmer: Jaccard** | |
| | **R2** | **p-val** | **R2** | **p-val** | **R2** | **p-val** | **R2** | **p-val** | **R2** | **p-val** | **R2** | **p-val** | **R2** | **p-val** | **R2** | **p-val** |
| **Canton** | 0.119 | **0.001** | 0.101 | **0.001** | 0.222 | **0.001** | 0.189 | **0.001** | 0.181 | **0.006** | 0.099 | **0.001** | 0.368 | **0.001** | 0.087 | 0.185 |
| **Fertilizer** | 0.060 | 0.116 | 0.062 | **0.030** | 0.062 | 0.119 | 0.059 | 0.212 | 0.085 | 0.104 | 0.087 | **0.001** | 0.093 | 0.091 | 0.098 | 0.099 |
| **Disease management** | 0.059 | 0.140 | 0.059 | 0.053 | 0.058 | 0.128 | 0.098 | **0.014** | 0.110 | 0.320 | 0.139 | **0.023** | 0.051 | 0.273 | 0.056 | 0.669 |
| **Farming system** | 0.053 | 0.363 | 0.055 | 0.143 | 0.052 | 0.215 | 0.052 | 0.283 | 0.041 | 0.502 | 0.065 | 0.164 | 0.034 | 0.445 | 0.067 | 0.430 |
| **Irrigation** | 0.050 | 0.496 | 0.049 | 0.632 | 0.047 | 0.278 | 0.039 | 0.586 | 0.039 | 0.558 | 0.060 | 0.476 | 0.053 | 0.263 | 0.038 | 0.968 |
| **Soil treatment** | 0.055 | 0.250 | 0.055 | 0.170 | 0.036 | 0.557 | 0.055 | 0.253 | 0.036 | 0.613 | 0.062 | 0.289 | 0.019 | 0.726 | 0.075 | 0.323 |
| **Canton : Farming system** | 0.053 | 0.356 | 0.055 | 0.182 | 0.059 | 0.119 | 0.050 | 0.367 | 0.164 | **0.020** | 0.070 | 0.068 | 0.073 | 0.162 | 0.063 | 0.533 |
| **Canton : Fertilizer** | 0.046 | 0.747 | 0.052 | 0.360 | 0.027 | 0.816 | 0.036 | 0.638 | | | | | | | | |
| **Canton : Irrigation** | 0.049 | 0.559 | 0.052 | 0.333 | 0.068 | 0.070 | 0.021 | 0.921 | 0.055 | 0.345 | 0.058 | 0.639 | 0.031 | 0.456 | 0.038 | 0.880 |
| **Canton : Soil treatment** | 0.052 | 0.394 | 0.054 | 0.185 | 0.044 | 0.322 | 0.041 | 0.529 | | | | | | | | |
| **Residuals** | 0.405 | | 0.406 | | 0.325 | | 0.360 | | 0.290 | | 0.360 | | 0.277 | | 0.478 | |

***Supplementary Table 6****: Nested, multivariate permutational analysis of variance (PERMANOVA) to assess correlations between fungal (ITS) and bacterial (16S) communities in soil with environmental variables, calculated per factor as well as nested by region (Average temperature is measured 10 cm below ground). (p-values < 0.05 are considered significant and highlighted in bold)*

| | **ITS** | | | | | | | | **16S** | | | | | | | |
|---|---|---|---|---|---|---|---|---|---|---|---|---|---|---|---|---|
| | **Bray Curtis** | | **Jaccard** | | **Kmer: Bray Curtis** | | **Kmer: Jaccard** | | **Bray Curtis** | | **Jaccard** | | **Kmer: Bray Curtis** | | **Kmer: Jaccard** | |
| | **R2** | **p-val** | **R2** | **p-val** | **R2** | **p-val** | **R2** | **p-val** | **R2** | **p-val** | **R2** | **p-val** | **R2** | **p-val** | **R2** | **p-val** |
| **Canton** | 0.120 | **0.001** | 0.101 | **0.001** | 0.222 | **0.001** | 0.189 | **0.001** | 0.185 | **0.006** | 0.095 | **0.001** | 0.368 | **0.002** | 0.087 | 0.162 |
| **Average Temperature** | 0.062 | 0.089 | 0.056 | 0.171 | 0.071 | 0.081 | 0.053 | 0.230 | 0.095 | 0.052 | 0.074 | **0.023** | 0.052 | 0.242 | 0.087 | 0.173 |
| **Altitude** | 0.060 | 0.109 | 0.059 | 0.072 | 0.062 | 0.142 | 0.074 | 0.075 | 0.043 | 0.413 | 0.075 | **0.015** | 0.030 | 0.443 | 0.065 | 0.416 |
| **Annual Rainfall** | 0.050 | 0.494 | 0.053 | 0.355 | 0.043 | 0.497 | 0.064 | 0.129 | 0.065 | 0.178 | 0.066 | 0.154 | 0.058 | 0.230 | 0.046 | 0.739 |
| **Canton : Annual Rainfall** | 0.045 | 0.822 | 0.050 | 0.673 | 0.035 | 0.682 | 0.067 | 0.100 | 0.038 | 0.512 | 0.062 | 0.367 | 0.023 | 0.574 | 0.054 | 0.604 |
| **Canton : Average Temperature** | 0.050 | 0.474 | 0.052 | 0.456 | 0.032 | 0.773 | 0.042 | 0.458 | 0.074 | 0.147 | 0.068 | 0.102 | 0.128 | 0.069 | 0.037 | 0.860 |
| **Canton : Altitude** | 0.058 | 0.169 | 0.055 | 0.190 | 0.053 | 0.272 | 0.039 | 0.544 | 0.139 | **0.006** | 0.074 | **0.005** | 0.047 | 0.278 | 0.111 | 0.158 |
| **Residuals** | 0.556 | | 0.574 | | 0.483 | | 0.472 | | 0.361 | | 0.485 | | 0.294 | | 0.513 | |

***Supplementary Table 7****: Nested, multivariate permutational analysis of variance (PERMANOVA) to assess correlations between fungal communities in berries with various vineyard management practices (Fertilizer application, Disease management: organic or synthetic fungicide application, Farming System: organic vs conventional, Irrigation, Soil treatment: cover crops vs bare soil), calculated per factor as well as nested by region. (p-values < 0.05 are considered significant and highlighted in bold)*

| | **Bray Curtis** | | **Jaccard** | | **Kmer: Bray Curtis** | | **Kmer: Jaccard** | |
|---|---|---|---|---|---|---|---|---|
| | **R2** | **p-val** | **R2** | **p-val** | **R2** | **p-val** | **R2** | **p-val** |
| **Canton** | 0.121 | **0.045** | 0.076 | **0.006** | 0.153 | 0.050 | 0.206 | **0.001** |
| **Fertilizer** | 0.077 | 0.148 | 0.069 | **0.021** | 0.123 | 0.071 | 0.080 | 0.064 |
| **Disease management** | 0.106 | 0.065 | 0.059 | 0.178 | 0.110 | 0.089 | 0.068 | 0.095 |
| **Farming system** | 0.045 | 0.453 | 0.061 | 0.106 | 0.027 | 0.557 | 0.069 | 0.110 |
| **Irrigation** | 0.024 | 0.873 | 0.049 | 0.748 | 0.023 | 0.641 | 0.024 | 0.853 |
| **Soil treatment** | 0.051 | 0.358 | 0.048 | 0.770 | 0.058 | 0.242 | 0.060 | 0.177 |
| **Canton : Farming system** | 0.049 | 0.403 | 0.048 | 0.799 | 0.077 | 0.170 | 0.037 | 0.538 |
| **Canton : Fertilizer** | 0.047 | 0.425 | 0.060 | 0.142 | 0.023 | 0.673 | 0.036 | 0.551 |
| **Canton : Irrigation** | 0.038 | 0.590 | 0.050 | 0.658 | 0.038 | 0.413 | 0.024 | 0.871 |
| **Canton : Soil treatment** | 0.042 | 0.484 | 0.054 | 0.443 | 0.021 | 0.695 | 0.064 | 0.118 |
| **Residuals** | 0.399 | | 0.425 | | 0.349 | | 0.331 | |

***Supplementary Table 8:*** *Nested, multivariate permutational analysis of variance (PERMANOVA) to assess correlations between fungal communities in berries with environmental variables, calculated per factor as well as nested by region. (Average temperature is measured 2 m above ground). (p-values < 0.05 are considered significant and highlighted in bold)*

| | **Bray Curtis** | | **Jaccard** | | **Kmer: Bray Curtis** | | **Kmer: Jaccard** | |
|---|---|---|---|---|---|---|---|---|
| | **R2** | **p-val** | **R2** | **p-val** | **R2** | **p-val** | **R2** | **p-val** |
| **Canton** | 0.121 | **0.021** | 0.078 | **0.004** | 0.153 | **0.036** | 0.206 | **0.001** |
| **Average Temperature** | 0.074 | 0.123 | 0.051 | 0.594 | 0.036 | 0.414 | 0.053 | 0.218 |
| **Altitude** | 0.049 | 0.301 | 0.062 | 0.097 | 0.053 | 0.252 | 0.059 | 0.159 |
| **Annual Rainfall** | 0.072 | 0.139 | 0.061 | 0.162 | 0.078 | 0.129 | 0.061 | 0.148 |
| **Canton : Annual Rainfall** | 0.054 | 0.281 | 0.061 | 0.138 | 0.034 | 0.430 | 0.037 | 0.529 |
| **Canton : Average Temperature** | 0.113 | **0.036** | 0.054 | 0.444 | 0.169 | **0.021** | 0.057 | 0.161 |
| **Canton : Altitude** | 0.042 | 0.403 | 0.047 | 0.817 | 0.048 | 0.270 | 0.071 | 0.085 |
| **Residuals** | 0.474 | | 0.586 | | 0.430 | | 0.456 | |

***Supplementary Table 9:*** *Permutational multivariate analysis of variance (PERMANOVA) of fungal beta diversity metrics in berry samples against berry chemistry parameters. (p-values < 0.05 are considered significant and highlighted in bold)*

| | **Bray Curtis** | | **Jaccard** | | **Kmer: Bray Curtis** | | **Kmer: Jaccard** | |
|---|---|---|---|---|---|---|---|---|
| | **R2** | **p-val** | **R2** | **p-val** | **R2** | **p-val** | **R2** | **p-val** |
| **Sugar content (degree Oechsle)** | 0.102 | 0.073 | 0.062 | 0.141 | 0.120 | 0.100 | 0.058 | 0.399 |
| **Assimilable nitrogen** | 0.079 | 0.180 | 0.062 | 0.135 | 0.096 | 0.141 | 0.043 | 0.643 |
| **Acid Total** | 0.027 | 0.874 | 0.059 | 0.246 | 0.025 | 0.673 | 0.088 | 0.113 |
| **Acid Tartaric** | 0.069 | 0.231 | 0.068 | 0.030 | 0.084 | 0.165 | 0.041 | 0.682 |
| **Acid Malic** | 0.036 | 0.714 | 0.045 | 0.911 | 0.028 | 0.593 | 0.052 | 0.502 |
| **pH berry** | 0.031 | 0.824 | 0.050 | 0.725 | 0.035 | 0.509 | 0.038 | 0.753 |
| **Residuals** | 0.656 | | 0.653 | | 0.613 | | 0.680 | |
